# Cryo-EM structures of Doravirine and Rilpivirine with HIV-1 Reverse Transcriptase/DNA Aptamer – Nonnucleoside Inhibitor Resistance by E138K and M184I Mutations

**DOI:** 10.1101/2022.02.27.482155

**Authors:** Abhimanyu K. Singh, Brent De Wijngaert, Marc Bijnens, Kris Uyttersprot, Hoai Nguyen, Sergio E. Martinez, Dominique Schols, Piet Herdewijn, Christophe Pannecouque, Eddy Arnold, Kalyan Das

## Abstract

Structures trapping a verity of functional and conformational states of HIV-1 reverse transcriptase (RT) have been determined by X-ray crystallography. These structures have played important roles in understanding the mechanisms of catalysis, inhibition and drug resistance, and in driving drug design. However, structures of several desired complexes of RT could not be obtained even after many crystallization or crystal soaking experiments. The ternary complexes of doravirine and rilpivirine with RT/DNA are such examples.

Structural study of HIV-1 RT by single-particle cryo-EM has been challenging due to the enzyme’s relatively smaller size and higher flexibility. We optimized a protocol for rapid structure determination of RT complexes by cryo-EM and determined six structures of wild-type and E138K/M184I mutant RT/DNA in complexes with the nonnucleoside inhibitors rilpivirine, doravirine, and nevirapine. RT/DNA/rilpivirine and RT/DNA/doravirine complexes have structural differences between them and from the representative conformation of RT/DNA/nevirapine (or efavirenz); the primer grip in the RT/DNA/doravirine and the YMDD motif in the RT/DNA/rilpivirine complexes have large shifts. The DNA primer 3’-end in the doravirine-bound structure is positioned at the active site, but the complex is in a non-productive state. In the mutant RT/DNA/rilpivirine structure, I184 is stacked with the DNA such that their relative positioning can influence rilpivirine in the pocket. Simultaneously, E138K mutation widens the NNRTI-binding pocket entrance, potentially contributing to a faster rate of rilpivirine dissociation by E138K/M184I mutant RT, as reported by earlier kinetic studies. These structural differences have implications for drug design and for understanding molecular mechanisms of drug resistance.

## Introduction

The emergence of drug resistance continues to be a challenge in long-term management of HIV-1 infection. Over four decades of extensive research on HIV and its modules has helped in improving the treatment options and quality of life for HIV-infected individuals. More than thirty drugs have been approved, and various drug combinations are used to treat the infected individuals under antiretroviral therapy (ART) regimens (1, 2); currently, over twenty-seven million individuals are taking antiviral drugs (WHO, 2021; https://www.who.int/news-room/factsheets/detail/hiv-drug-resistance). The individual drugs target specific functional steps in the lifecycle of the virus, such as viral attachment and fusion, reverse transcription, integration of viral DNA to the host-cell DNA, and maturation of newly formed immature virions.(3, 4) A major challenge in prolonged antiviral treatment is the emergence of drug-resistant mutant strains. The mutations that confer resistance to a drug commonly appear in the targeted protein, and often the mutations directly or indirectly afflict the binding of a drug.

The viral enzyme reverse transcriptase (RT) copies the viral single-stranded RNA (ssRNA) to a double-stranded DNA. RT is targeted by thirteen drugs of which seven are nucleoside RT inhibitors (NRTIs) (5) and six are non-nucleoside RT inhibitors (NNRTIs) – nevirapine (NVP), delavirdine, efavirenz (EFV), etravirine, rilpivirine (RPV), and doravirine (DOR). RT mutations confer resistance to NRTIs and NNRTIs. (6-9) Biochemical and structural studies help in understanding the molecular mechanisms of drug-resistance mutations. NRTIs are generally DNA chain terminators. The dNTP-binding site and surrounding amino acid residues are mutated to confer resistance to NRTIs. The primary mechanisms of NRTI resistance mutations are steric hindrance such as by M184V/I, discrimination by K65R, and excision of AZT.(10, 11) NNRTIs bind to a hydrophobic pocket known as non-nucleoside inhibitor binding pocket (NNIBP), which is located adjacent to the polymerase active site. The pocket mutations cause NNRTI resistance primarily by steric hindrance and loss of inhibitor-protein interactions. Biochemically, mutations at the NNIBP entrance are shown to develop NNRTI resistance by increasing the rate of NNRTI dissociation from NNIBP. Crystal structures and structure-based approaches have aided the design of new NNRTIs. (12-14)

Structures of HIV-1 RT trapping various functional and conformational states have been determined by X-ray crystallography. (15) However, structures of several desired complexes of RT could not be achieved when crystallization or crystal soaking experiments were not successful. The ternary complexes of DOR and RPV with RT/DNA are such examples. The clinically emerging mutations are different for different NNRTIs. (16) (https://hivdb.stanford.edu/dr-summary/resistance-notes/NNRTI/) The predominant clinical mutations associated with resistance to rilpivirine are E138K/M184I(or V). (17, 18) The ternary structure of wild-type RT/DNA/RPV and (E138K/M184I) RT/DNA/RPV can help unfold the roles of these two mutations that are primarily outside the NNIBP and far apart from each other. Our study focuses on finding the molecular mechanism of RPV resistance and visualizing why another second-generation NNRTI DOR is not impacted by the double mutation.

RT is a heterodimer of p66 and p51 subunits with a combined molecular mass of 117 kDa. RT is composed of multiple subdomains, and the movement of the subdomains, which is essential for carrying out its functions, makes RT highly flexible in solution. Because of its relatively smaller size and higher flexibility, obtaining atomic-resolution structures of RT and its complexes by single-particle cryo-electron microscopy (cryo-EM) has been challenging. A recent study reported structures of RT/dsRNA/NNRTI (NVP and EFV) complexes at resolutions ranging between 2.8–3.1 Å. (19) A large amount of data was acquired and processed to get those structures from final sets of 1.2 million-plus particles. In contrast, a high-throughput experimental platform is essential for the effective use of the single-particle cryo-EM technique for large-scale structure determination of various RT complexes and for using structures in drug design. In the current study, we are reporting the structures of wild-type and E138K/M184I mutant RT/DNA aptamer (DNA-apt) in complexes with RPV, DOR, and NVP; the resistance mutations E138K is in p51 and M184I is in the p66 subunit, respectively. (20) The structures show how the positioning of the DNA with respect to I184 and opening of the NNIBP entrance by E138K mutation contribute to drug resistance.

## Results

### High-throughput cryo-EM structures of RT/DNA/NNRTI complexes

We have determined six structures of wild-type HIV-1 RT/DNA and M184I/E138K mutant RT/DNA in complexes with the NNRTIs RPV, DOR, and NVP at resolutions ranging from 3.32 to

3.65 Å by single-particle cryo-EM (SI Table 1). Recently, we trapped an intermediate P-1 complex of RT and determined the cryo-EM structures of the complex with small-molecule fragments bound at a transient P pocket. (21) Building on those experiments, we optimized a cryo-EM platform for routine and rapid structure determination of catalytically active RT/DNA and inhibited RT/DNA/NNRTI complexes.

A 38-mer DNA aptamer (DNA-apt; Fig. 1A) binds HIV-1 RT with ∼15 pM affinity, which is ∼100x higher than a regular dsDNA (22), and was successfully used for structural studies of RT/DNA-apt/dNTP (or analogs) by X-ray crystallography. (23, 24) In the current study, RT/DNA-apt complexes were purified over a size-exclusion column and ascertained as a homogenous complex by multi-angle light scattering (MALS) and dynamic light scattering (DLS) measurements (Fig. 1B). The pure and homogenous RT/DNA samples were incubated with RPV, DOR, or NVP to form the respective RT/DNA/NNRTI ternary complexes. The homogeneity of each ternary complex was tested by DLS, and highly reproducible vitreous grids of individual complexes were prepared on Quantifoil holey carbon Au or UltrAuFoil grids using a low sample concentration of ∼0.3 mg/ml. All datasets were collected on a Glacios 200 kV transmission electron microscope (TEM)/Falcon 3 setup as installed in our lab. In most cases, fewer than 2000 micrographs were recorded per complex of which a high percentage were of excellent quality with little to no contamination. The clean micrographs enabled picking good starting sets of particles that steered faster processing to high-quality density maps for each structure (Figs. 1C, D) including for the bound NNRTIs (Figs. 1E, F). This optimized protocol, as detailed in the Materials and Methods section, would enable routine use of cryo-EM for high-throughput structural study and structure-based drug design.

**Figure 1.**
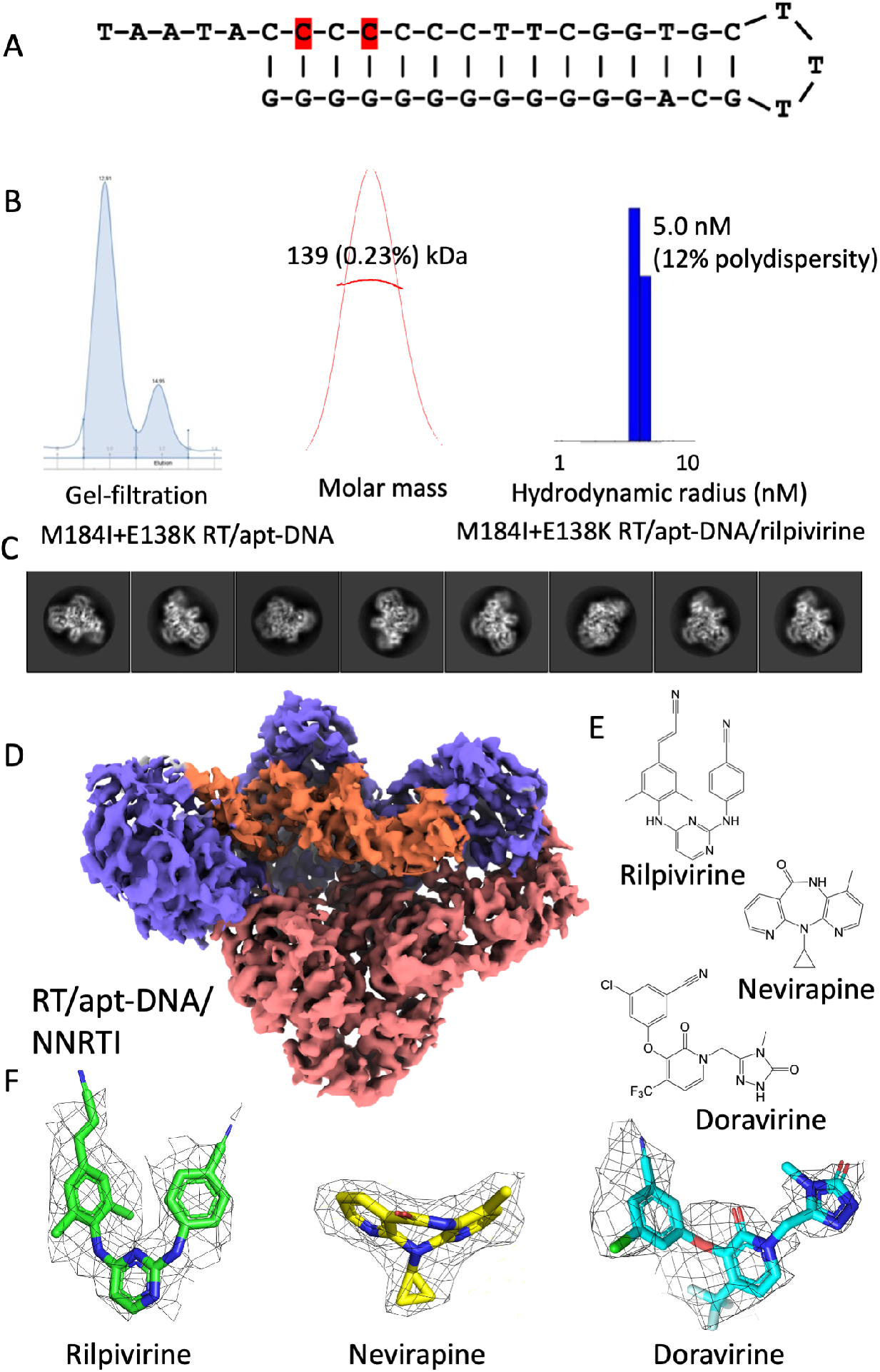
Sample and grid optimization for rapid cryo-EM structure determination. **A**. The DNA-aptamer used in this study to form RT/DNA complex; the aptamer in the wild-type RT/DNA/NNRTI complexes and the mutant RT/DNA/NNRTI complexes have a dTMP and dATP as the first template overhang, respectively. The modified 2-O-methyl dCMP nucleotides at the −2 and −4 positions are highlighted. **B**. RT/DNA complex purified by size-exclusion chromatography; MALS and DLS measurements ascertained the high quality of the sample for single-particle cryo-EM studies. **C**. Selected 2D classes symbolize the high-resolution characteristics of the complexes. **D**. Cryo-EM density map covering the RT/DNA/rilpivirine (RPV) complex, representing the RT/DNA/NNRTI ternary complex structures. **E**. Chemical structures of the NNRTI drugs rilpivirine (RPV), nevirapine (NVP), and doravirine (DOR) used in the current study. **F**. The segments of cryo-EM density maps covering the NNRTI drugs RPV, NVP, and DOR; the maps for RPV, NVP, and DOR are contoured at 3.2, 4, and 2.20, respectively. The stereo views of NNRTI fitted to the experimental density maps for all six structures are shown in SI Fig. 1.

### Structures of RPV, NVP, and DOR in complex with wild-type RT/DNA aptamer

The first structure determined in the series is of wild-type (wt) RT/DNA-apt/NVP which is analogous to the earlier reported crystal structures of RT/DNA/NVP (25) and RT/DNA/EFV (26). Experimentally, a DNA aptamer is used in the current cryo-EM study of the RT/DNA/NVP complex whereas, RT and DNA were cross-linked in the crystal structure. Both structures align well with RMSD of ∼0.7 Å for 757 superimposed Cα atoms (SI Fig. 2); NVP, NNIBP residues, and the cross-linked DNA and aptamer DNA align almost indistinguishably between two structures. The structure comparison shows that using either of two distinct structure determination techniques (X-ray and cryo-EM), with either DNA cross-linking or a DNA aptamer, all lead to a very consistent structure of the RT/DNA/NVP complex. For simplification, hereafter, we refer DNA-apt as DNA throughout the text except where both are directly compared.

The density map for the wild-type RT/DNA/RPV ternary complex (Fig. 2A) was obtained at 3.38 Å resolution. The ternary complex superimposes on the crystal structure of the RT/RPV binary complex (27) with RMSD of 1.5 Å for 876 Cα atoms. The conformation and the mode of binding of RPV however are not significantly impacted by the binding of DNA (Figs. 2B-C). An important structural difference between the binary and ternary complexes of RPV is that upon DNA binding, the active-site YMDD motif is lifted towards the DNA by about 2.2 Å at the M184 position.

**Figure 2.**
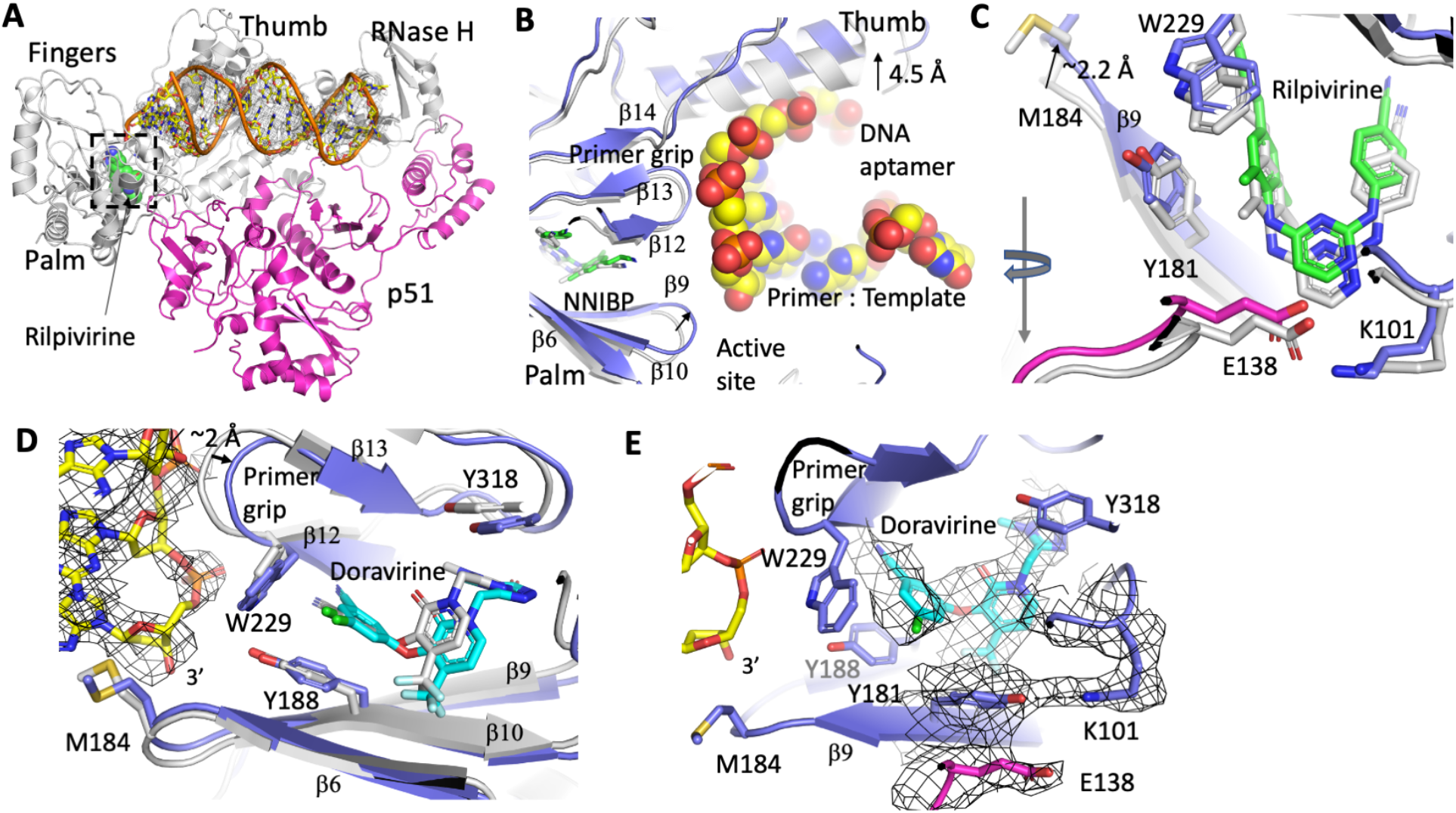
Structural changes upon DNA binding to RT/RPV and RT/ DOR complexes. **A**. Structure of wild-type RT/DNA/RPV complex; density for the DNA is displayed at 2.50. **B**. A zoomed view of the polymerase active site showing the Cα-superposition of RT/RPV binary complex (PDB Id. 4G1Q; gray) and RT/DNA/RPV ternary complex structures (blue protein, space-filled DNA, and green RPV); 876 Cα atoms superimposed with RMSD of 1.5 Å. There are subtle but important differences between two structures. The tip of the thumb moves outward by ∼4.5 Å to accommodate DNA. **C**. A zoomed view of the NNIBP region of the superimposed binary and ternary complexes. The YM_184_DD motif has moved up by ∼2 Å in the ternary complex structure. The inhibitor protein interaction and E138 … K101 salt-bridge link are conserved in both structures. **D**. Cα-superposition of RT/DOR binary (PDB Id. 4NCG; gray) and the RT/DNA/DOR ternary complex structures (blue protein, yellow DNA, and cyan DOR); 793 Cα atoms superimposed with RMSD of 0.99 Å. The binding of DNA to RT/DOR repositions the primer grip. **E**. Cryo-EM density defines the binding of DOR in the RT/DNA/DOR ternary complex. Unlike in the RPV ternary structure, E138 of p51 stacks with the Y181 sidechain, which forms a hydrogen bond with K101; the density map is contoured at 2.2σ.

The cryo-EM structure of the RT/DNA/DOR ternary complex was determined at 3.65 Å resolution. The structure superimposes on the RT/DOR binary complex (28) with RMSD of 0.99 Å for 793 Cα atoms. The binding of DNA has a significant impact on the primer grip-containing β12−β13−β14 sheet; the primer grip is pushed by about 2 Å to accommodate the DNA (Fig. 2D). In contrast, the β6−β10−β9 sheet, that contains the YMDD motif, is less perturbed by the DNA binding; the residue Y181 has an altered rotamer in both RT/DOR and RT/DNA-apt/DOR structures (Fig. 2E) when compared to that in other structures. (Fig. 2C) The change in NNIBP due to a large shift of β12−β13−β14 sheet is non-uniform, and consequently, DOR is rearranged by repositioning and readjusting its rotatable bonds (Fig. 2D) to adapt to the pocket changes upon DNA binding. In general, the comparisons of ternary and binary complexes of NNRTIs show that DNA binding has different impacts on different NNRTIs. Structurally, RPV and DOR, each has five rotatable bonds providing torsional freedoms which help the NNRTIs wiggle and jiggle to adapt to changes in NNIBP. (14)

### DOR and RPV ternary complex deviate from the standard RT/DNA/NNRTI conformation

Superposition of RT/DNA/NNRTI ternary complex structures shows that the p66 connection, RNase H, and the p51 subunit form a structurally stable core. The polymerase domain, which is composed of fingers, palm, and thumb, has positional differences among the structures. (Fig. 3A) These differences are rather non-uniform and influenced by the bound NNRTIs. The key structural elements (i) β12−β13−β14 sheet that contains the primer grip and(ii) β6−β10−β9 sheet that contains the polymerase active site are known to be involved in the binding of nucleic acid substrates (dsRNA, dsDNA, and RNA/DNA) as well as the NNRTIs. These structural elements align well in the RT/dsRNA mini-transcription initiation ternary complexes with or without NNRTIs (19), RT/DNA/NVP (25), and RT/RNA-DNA/EFV (26). In fact, the NNRTI-bound conformation of RT has a close resemblance with RT/dsRNA transcription initiation state (SI Fig 3); only the track of dsRNA is different from that of dsDNA. (29) Therefore, the positioning of the primer grip, YMDD motif, and DNA in the RT/DNA/DOR and RT/DNA/RPV complexes were expected to align with those in the RT/DNA/NVP complex. Surprisingly, the DOR and RPV bound RT/DNA ternary complex structures have significant perturbation from the standard conformation of the RT/DNA/NNRTI structures; the positioning of the primer grip and the YMDD motif have considerable deviations in the DOR and RPV ternary complexes, respectively. (Fig. 3B) Interestingly, in the RT/DNA/DOR structure, the primer 3’-end is positioned near the active site, in contrast to >5 Å shift of the primer 3’-end in all other RT/DNA/NNRTI structures including RT/DNA/RPV. (Figs. 3C and SI Fig. 3B)

**Figure 3.**
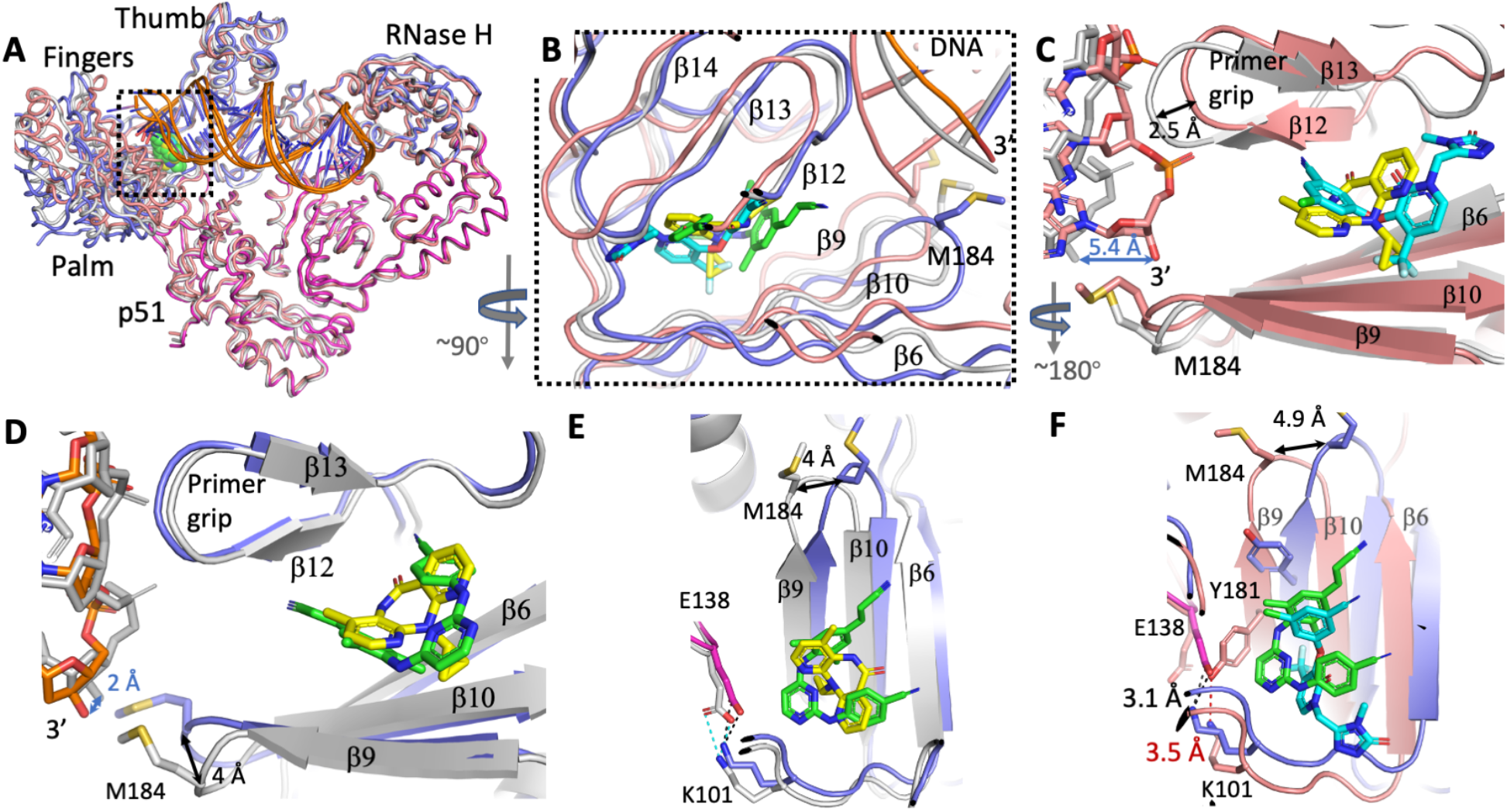
RT/DNA/RPV and RT/DNA/DOR structures deviate from the standard conformation of RT/DNA/NNRTI complexes represented by RT/DNA/NVP. **A**. Cα-superposition of RT/DNA/RPV (blue p66, magenta p51), RT/DNA/DOR (salmon) and RT/DNA/NVP (gray). The p51 and p66 connection, and RNase H form a structurally stable core; large structural variations are observed for the polymerase domain consists of fingers, palm, and thumb. A superpositions of these structures with RT/dsRNA/NNRTI complexes are shown in SI Fig. 3. **B**. A zoomed view of the superimposed structures shows repositioning of the β6−β10−β9 sheet that contains the YMDD motif and β12−β13−β14 sheet that contains the primer grip; the RT/DNA and NNRTIs are colored blue and green in the RPV complex, salmon and cyan in the DOR complex, and gray and yellow in the NVP complex, respectively. **C**. Superimposed RT/DNA (salmon)/DOR (cyan) and RT/DNA (gray)/NVP (yellow) structures show that the DNA primer 3’-end has moved by ∼5.4 Å towards the active site in the DOR complex compared to that in NVP complex. Consequently, the primer grip is shifted by ∼2.5 Å in the DOR complex to accommodate the repositioned DNA. **D**. Superimposed RT (blue)/DNA (orange)/RPV (green) and RT/DNA (gray)/NVP (yellow) structures show a common DNA track; however, the YMDD motif has shifted up by ∼4 Å in the RPV complex. **E**. A view of the superimposed RPV and NVP structures down the primer grip shows a sliding motion for the β6−β10−β9 sheet upon RPV binding. The K101 … E138 salt-bridge at about the 7 o’clock position is conserved in both structures; E138 in RPV complex is in magenta. **F**. The superposition of RTs in RPV (blue p66 and magenta p51 of RT and green RPV) and DOR (salmon RT and cyan DOR) ternary complexes also shows the sliding motion of the β6−β10−β9 sheet in response to the binding of DNA and RPV compared to DOR. The altered rotamer of Y181 in the DOR complex forms an H-bond with K101 at the NNIBP entrance, and E138 switched its conformation to stack with Y181; this stacking of E138 with Y181 is also present in the RT/DOR binary complex structure.

Current RPV and NVP ternary complex structures superimpose with 0.75 Å RMSD for 757 Cα atoms (Fig. 3D); the DNAs follow a common track, and the primer grips align, however, the YMDD motifs are positioned differently at ∼4 Å apart. The YMDD-containing β6−β10−β9 sheet in RT/DNA/RPV has skidded significantly from its position in the NVP and DOR ternary complexes. (Figs. 3E, F) The DNAs in RPV and NVP structures have common, yet not highly superimposable tracks; the 3’-end nucleotides in RPV and NVP ternary complex structures are ∼ 2 Å apart. Like in most RT/NNRTI crystal structures, the p51 E138 side chain forms a salt bridge with the p66 K101 side chain at the NNIBP entrance in both ternary structures, however, RPV is positioned closer to the NNIBP entrance with a minimum distance of ∼3.5 Å from E138 whereas, NVP is positioned deeper in the pocket at a minimum distance of ∼5.4 Å from E138. (Fig. 3E) In the DOR ternary complex, the switched conformation of Y181 forms an H-bond with K101 and the E138 side chain stacks with Y181. (Figs. 2E & 3F) The structures reveal that the binding of second-generation NNRTIs (RPV or DOR) and DNA influence one another, as well as positioning of the key structural elements of RT. These structural differences appear to have roles in developing NNRTI resistance.

### The DNA track is less perturbed by DOR than by other NNRTIs

Biochemically, NNRTIs are known to inhibit the catalytic incorporation of nucleotides. (30, 31) The distortion of the catalytic triad or arrest of the primer grip, or most likely the combination of both contribute to the NNRTI inhibition. An earlier structural study also revealed that NVP binding impedes the binding of dNTP. (25) Surprisingly, the DNA track in the DOR complex aligns with that of the catalytically active RT/DNA-apt structure (SI Fig. 4A) (23), which is very different from that in all other RT/DNA/NNRTI ternary complex structures (SI Figs. 4B-C), indicating the possibility that the mechanism of inhibition may be somewhat different for DOR. The possible modes of inhibition by DOR are (i) like for other NNRTIs, RT/DNA/DOR complex weakens the binding of dNTP at the polymerase active site, (ii) an incoming dNTP is bound however, DOR blocks its catalytic incorporation, or (iii) one nucleotide is incorporated and the following translocation step is blocked by DOR jamming the primer grip. To investigate the precise mode of DOR inhibition, we attempted the cryo-EM structure of RT/DNA/DOR in the presence of ∼100x molar excess of dATP as the incoming dNTP. The structure showed no density for dATP at the active site. A dNTP-binding is associated with the fingers closing (32); however, the fingers in our structure remained open confirming no dTTP binding. Our single-nucleotide incorporation assay (SI Fig. 5), as discussed later, showed no nucleotide incorporation in the presence of DOR. These results confirm that DOR inhibits RT in an analogous fashion as other NNRTIs despite the DNA primer terminal being positioned at the active site in the RT/DNA/DOR complex, i.e., the DNA binds in a non-productive state in the presence of DOR despite its 3’-end is positioned at the polymerase active site. The track of DNA in the RT/DNA/DOR complex is guided by the positioning of the primer grip, which is different from that in all other structures. (Fig. 3C & SI Fig. 4) Consequently, the P-1 nucleotide of the DNA that interacts with the primer grip has dipped by ∼ 4 Å with the primer grip to position the 3’-end at the active site.

### Structures of E138K/M184I mutant RT/DNA in complexes with RPV, NVP, and DOR

We determined the cryo-EM structures of E138K/M184I mutant RT/DNA in complexes with NVP, RPV, and DOR at 3.38, 3.45, 3.58 Å resolution, respectively (SI Table 1). The (E138K/M184I) RT/DNA/NNRTI structures align well with their respective wild-type RT/DNA/NNRTI structures; 879 Cα atoms align with 0.62 Å RMSD for the RPV complex, 906 Cα atoms aligned with 0.69 Å RMSD for DOR complexes, and 838 Cα atoms aligned with 0.64 Å RMSD for NVP complexes. Two mutation sites, E138K at the entrance to NNIBP and M184I of YMDD motif, are ∼15 Å apart in RPV-, ∼13 Å in NVP-, and ∼11 Å in DOR-bound ternary complexes; this distance is ∼11 Å in the active RT/DNA complex with no bound NNRTI. An earlier structural study had shown that the loss of a salt bridge between E138 and K101 in the E138K mutant is the primary contributor to NNRTI resistance. (33) The current cryo-EM density maps of wild-type and mutant RT/DNA/NVP and RT/DNA/RPV complexes show the loss of the salt bridge and widening of the NNIBP are caused by the E138K mutation (Figs. 4A & B); RPV is positioned closer to the NNIBP entrance. The entrance in the DOR-bound structure is least influenced by the E138K mutation; the aromatic side chain of Y181 that blocks the entrance (Fig. 3F) in the wild-type complex loses the H-bond with K101 in the mutant complex, however, the Y181 sidechain still blocks the NNIBP entrance which may prevent faster dissociation of DOR from NNIBP (Fig. 4C).

**Figure 4.**
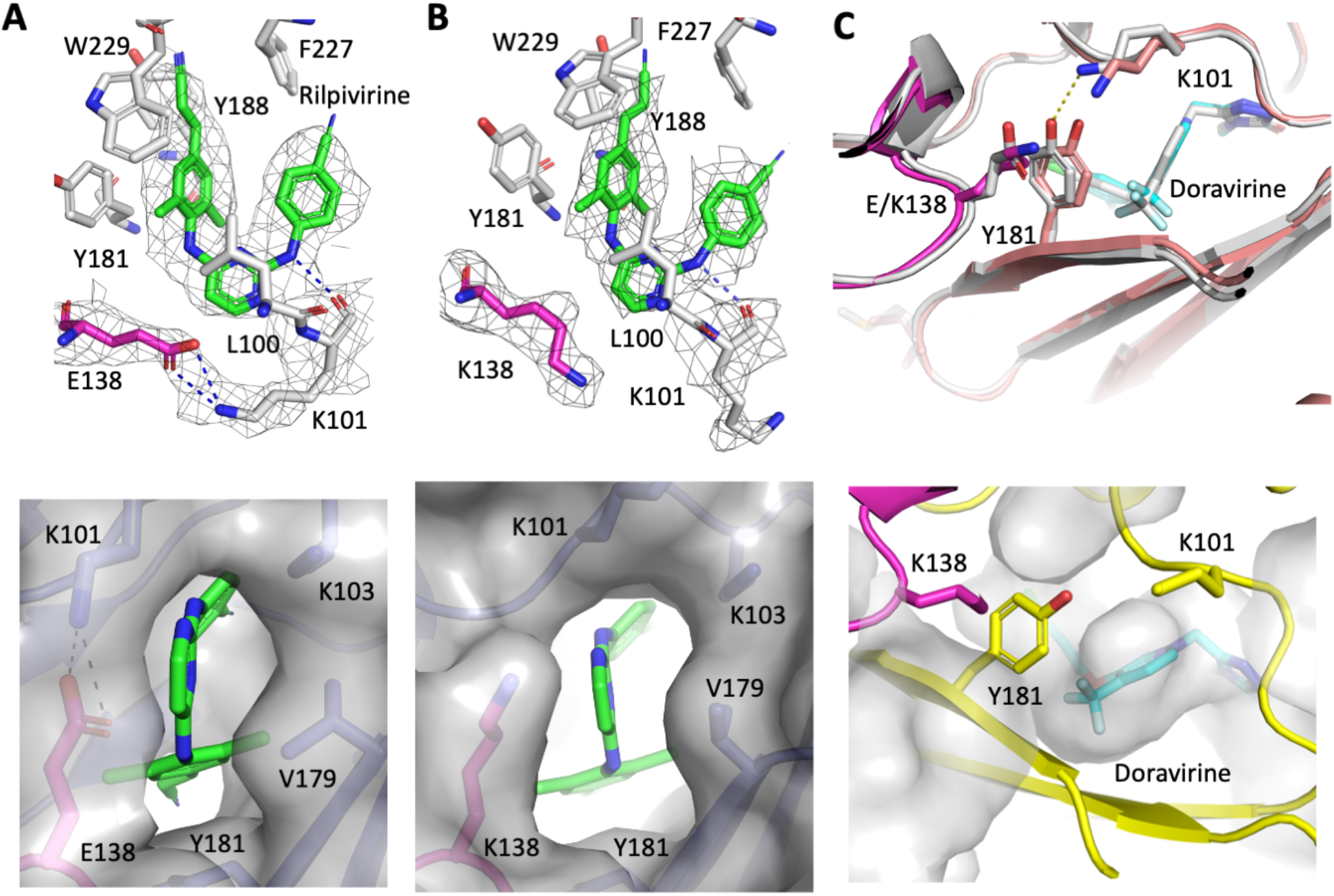
E138K mutation at the NNIBP entrance in RPV and DOR ternary complexes. RPV and surrounding residues in wild-type (**A**) and M184I/E138K RT (**B**) ternary complexes. The bottom panels show the molecular surfaces at the NNIBP entrance of respective structures. The cryo-EM density is shown for RPV, E/K138 (p51) and K101. The loss of the H-bond between E138 and K101 by E138K mutation widens the entrance to the pocket. **C**. A comparison of the pocket entrances in the wild-type and (E138K/M184I) mutant RT/DNA/DOR complexes reveal no significant impact of the mutations at the NNIBP entrance; the entrance is covered by the Y181 side chain in both RT/DNA/DOR structures. The molecular surface of DOR in the bottom panel gives an impression of the entrance blocked by Y181.

M184I/V is an NRTI-resistant mutation that reduces the susceptibility of L-nucleotide analogs lamivudine (3TC) and emtricitabine (FTC) to HIV (34, 35). Kinetic studies and crystal structures have revealed that the 3TCTP (or FTCTP) binds M184I/V RT in a non-productive mode with lower affinity when compared to dCTP. (32, 36, 37) However, it was not clear how M184I/V mutation contributes to NNRTI resistance. Both E138K and M184I mutations have mild impacts on NNIBP residues, the track of DNA, and the positioning of YM(I)DD loop in the RPV-bound structures (Figs. 5 A & B). The cyanovinyl group attached to the dimethylphenyl ring of RPV has flipped in the mutant structure when compared to the wild-type complex; the experimental density, and relative positioning of the dimethylphenyl ring and surrounding residues confirm the flipped orientation of the cyanovinyl group. The adaptability of the cyanovinyl group and its interactions with RT are critical for the binding of RPV and for retaining its potency against several NNRTI-resistant mutations. (14, 27) The relative positioning of the I184 side chain with respect to the DNA primer 3’-end nucleotide is distinct in the RPV complex when compared to that in the mutant RT/DNA/DOR (Fig. 5D) and RT/DNA/NVP (Fig. 5E) structures. The I184 side-chain and the sugar-phosphate backbone of the 3’-end nucleotide are stacked (Fig. 5F). It is clear from the structures that in a dynamic state in which RT slides and flips over a nucleic acid substrate, (38) the YM(I)DD motif in the RPV complex would respond differently than that in other NNRTI ternary complexes. In the RPV complex, I184 and the DNA face each other, which may restrict the motion of YM(I)DD loop, whereas in the NVP and DOR complexes, the YM(I)DD motif would be able to move up and down like a springboard, which is also the natural motion of the loop for carrying out DNA polymerization. The ability of RPV to wiggle and jiggle (14) would help compensate for the potential impact from M184I mutation alone, however, a wider NNIBP entrance in E138K-mutant RT would help in increasing the rate of dissociation of RPV. These observations support the earlier biochemical studies which showed that viruses containing only I184 would retain full susceptibility towards RPV whereas, the E138K/M184I mutant reduces RPV susceptibility by lowering RPV’s equilibrium binding affinity to the mutant RT. (39, 40)

**Figure 5.**
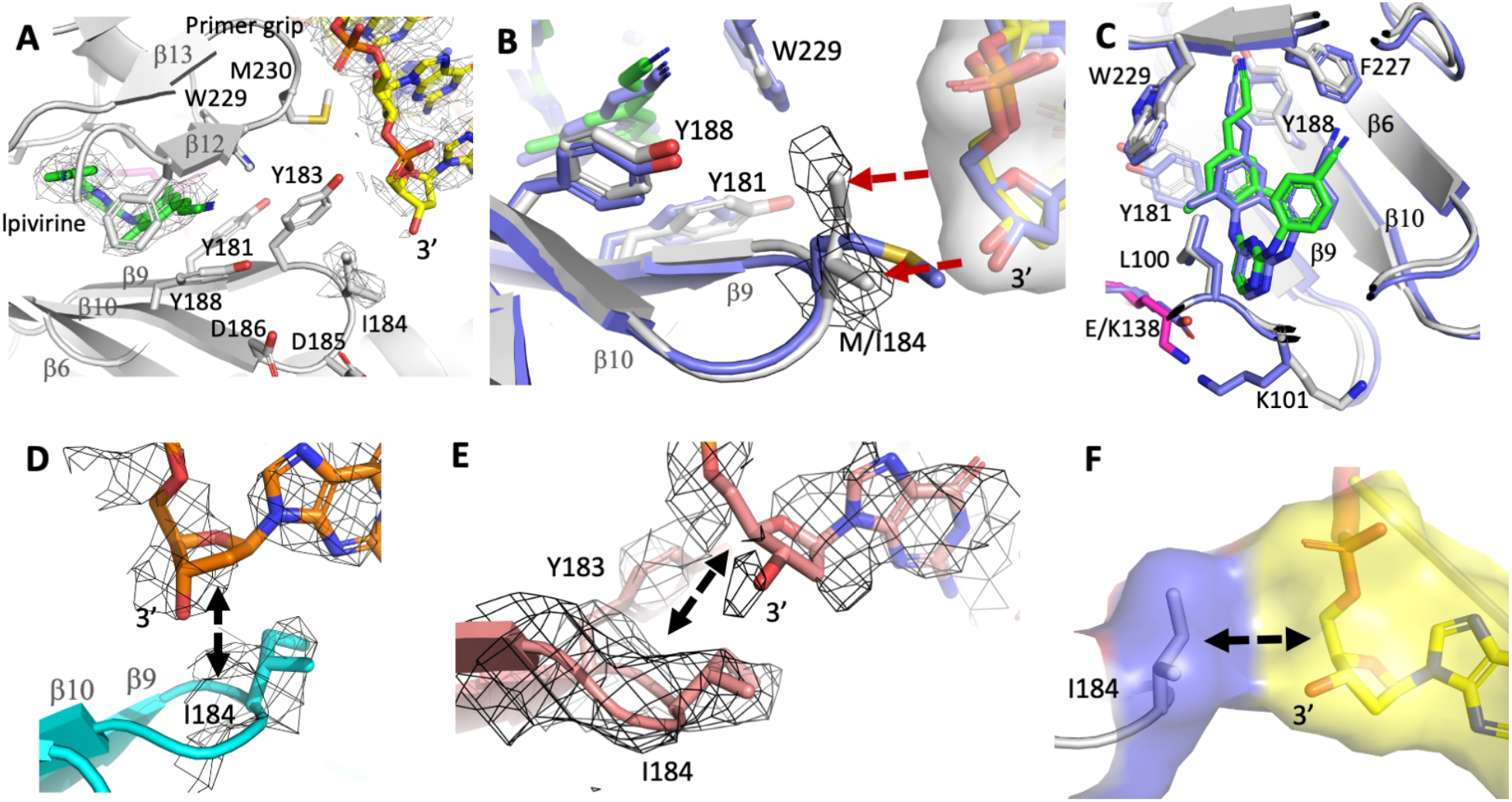
Relative positioning of the mutated I184 residue with respect to DNA primer 3’-end is distinct in the RPV-bound complex when compared to DOR and NVP ternary complexes. **A**. The NNIBP and polymerase active site in the (M184I/E138K) RT (gray)/DNA (yellow)/RPV (green) structure; the cryo-EM density for I184 and DNA is contoured at 2.00. **B**. Superposition of the wild-type RT/DNA/RPV structure (blue) on the mutant RT (gray)/DNA (yellow)/RPV (green) complex shows that the β-branched rigid side chain of I184 compared to a more flexible M184 side chain is locked against the DNA primer 3’-end nucleotide. **C**. The binding mode of RPV (green) in the mutant RT (gray)/DNA (yellow) ternary complex has its cyanovinyl group flipped when compared to the wild-type RT/DNA/RPV (blue) complex. **D**. Relative positioning of the DNA primer 3’-end with respect to the YM(I)DD motif in the (M184I/E138K) RT (cyan)/DNA (orange)/DOR structure. **E**. Relative positioning of the DNA primer end with respect to the YM(I)DD motif in the (M184I + E138K) RT/DNA (salmon)/NVP structure. The displayed sections of the cryo-EM densities in panels D and E are contoured at 2.00. **F**. The molecular surface showing the extensive interaction between I184 and the backbone of the 3’-end nucleotide in the (M184I/E138K) RT/DNA/RPV ternary complex.

### E138K and M184I mutations in clinical and biochemical context

E138K is a non-polymorphic mutation selected in patients on RPV-containing treatment regimens. It frequently emerges in combination with M184I in patients experiencing virological failure. (18) Although it is difficult to ascertain which of these two mutations emerges first, it has been suggested that the E138K mutation is likely selected for RPV if no other mutation is present. (41) Following the identification of E138K and M184V/I as characteristic RPV-resistance mutations in clinic, studies have been conducted to see the effects of the mutations in cell-based assays and at the enzyme level. Enzymatically, E138K/M184I RT has a moderate increase in resistance to RPV (40, 42). Phenotypic assays have shown that E138K in the p51 subunit alone decreases RPV susceptibility by about 2.4-fold while the co-existence of M184I in p66 further brings the susceptibility down to ∼5-fold (39); M184I/V mutation alone, however, does not alter RPV susceptibility. M184I/V is a nucleoside-resistance mutation, which negatively affects the RT replication capacity as shown earlier by steady-state kinetic experiments; Km of 14.6 for the mutant vs. 5.6 µM for the wild-type enzyme. (43) The decrease in enzyme functionality, however, gets compensated by E138K in p51 which restores the K_m_ for dNTP to a value close to the wild-type enzyme. Thus, the clinical selection of E138K and M184I mutations seems to play a dual role: conferring RPV resistance and maintaining the viral fitness level. The effect of E138K mutation on DOR susceptibility has been also studied. E138K has not been selected against DOR in clinic or shown to affect DOR susceptibility to a noticeable degree *in vitro*. (44, 45) DOR and RPV are shown to have complementary resistance mutation profiles. (46)

To investigate the inhibitory effects of NVP, RPV, and DOR on the wild-type and E138K/M184I RTs, we employed two separate sets of biochemical assays. Inhibition assay to determine IC_50_ values (50% inhibitory concentration) for the inhibitors on the wild-type and (E138K/M184I) RTs were carried out using EnzChek™ Reverse Transcriptase Assay Kit (ThermoFisher Scientific) (Table 1). Our assay shows that NVP inhibits both wild-type and the mutant RT at a comparable IC_50_; however, reduced susceptibility for NVP by the E138K/M184I mutant RT of HIV-1 subtype C has been previously shown in a phenotypic assay. (9) For both RPV and DOR, our assay results show a moderate decrease in susceptibility by E138K/M184I RT over the wild-type RT with a resistance fold change of ∼2.3 and 2.4, respectively (Table 1).

**Table 1:** Inhibitory activity of NVP, RPV and DOR on wild-type and E138K/M184I RT.

| IC<sub>50</sub> (nM) |  |  |  |
| --- | --- | --- | --- |
| RT | NVP | RPV | DOR |
| Wild-type | 130 ± 10 | 3.9 ± 0.4 | 11 ± 1 |
| E138K/M184I | 88 ± 17 (0.7)<sup>#</sup> | 8.9 ± 0 (2.3) | 26 ± 1 (2.4) |

<sup>#</sup> Value in parentheses represents resistance fold change with respect to the wild-type RT. Standard deviation for each measurement is indicated.

In parallel, we have also performed a gel-based single-nucleotide incorporation assay (47) to assess the effect of NVP, RPV, and DOR on DNA polymerization by RT. All three NNRTIs inhibit DNA polymerization by wild-type and E138K/M184I RTs in a dose-dependent manner with the mutant RT showing slightly reduced susceptibility towards all three inhibitors (SI Fig. 5).

## Discussion

The structural adaptability of the second-generation NNRTIs such as RPV and DOR helps the drugs overcome the impacts of common NNRTI-resistance mutations such as Y181C, which is deleterious to NVP. The binding pocket is predominantly hydrophobic, and so are the NNRTIs. Biochemically, the impacts of E138K or E138K/M184I mutations on NNRTIs are rather low in our assays which are in agreement with prior studies; however, the mutations predominantly emerge in patients in RPV regimen. The current cryo-EM structures of the NNRTI ternary complexes show that a wider pocket entrance and the relative positioning of I184 with respect to the DNA substrate in the RPV-bound structure appear to contribute to RPV resistance by E138K/M184I mutations. In contrast, the track of DNA is less influenced by the binding of DOR, and the NNIBP entrance is almost unaltered by p51 E138K mutation. The size and shape of the NNRTIs influence the arrangement of the key structural elements – YMDD motif, primer grip, track of DNA, the pocket entrance, etc. The relative distance between the two mutation sites E138K (p51) and M184I (p66) also vary among RT/DNA/NNRTI ternary complex structures; the distance between the two Cα positions is maximum in RPV complex whereas, the distance is not altered in DOR complex when compared to polymerase active RT/DNA complex. This result suggests that large structural changes to the RT/DNA elongation complex induced by the binding of an NNRTI may have a structural disadvantage that is susceptible to resistance mutations. Therefore, it may worth routine analysis of the structural features of NNRTI-inhibited RT/DNA (or RNA/DNA) elongation complexes for effective design of NNRTIs. The optimized single-particle cryo-EM protocol outlined in this study will help overcome the limitation of growing crystals of desirable wild-type and mutant HIV-1 RT/DNA/NNRTI complexes. The single-particle cryo-EM technique has been very powerful in obtaining new structures of challenging macromolecule complexes in recent years. The current study provides one of the few examples of how the cryo-EM technique can be optimized for rapid structure determination to aid drug design.

## Materials and Methods

### RT double-mutant construction

To create the HIV-1 RT double mutant containing E138K and M184I mutations in p51 and p66 domains respectively, two separate pETDuet-1 vectors harboring p51 and p66 coding segments were used. The two domains constitute an RT construct (RT139A) as reported previously. (48) Overlapping primers (p51-For-E138K-ATTAACAATAAAACTCCCGGGATC and p51-Rev-E138K-GGAGTTTTATTGTTAATGCTAGGGAT; p66-For-M184I-ATTTATCAATATATAGATGACTTGT and p66-

Rev-M184I-CATACAAGTCATCTATATATTGA) were used in PCR to introduce the desired nucleotide changes. One µl of each plasmid at ∼1.5 ng/µl concentration were used as template and the reactions were carried out with Q5 High-Fidelity DNA Polymerase (New England Biolabs). PCR products were gel purified and successful mutagenesis was confirmed by sequencing (Macrogen). The amplified plasmid containing the mutated p66 was double digested with *Nde*I and *Xho*I restriction enzymes (New England Biolabs), gel purified and cloned into MCS-2 of the pETDuet-1 vector pre-digested with the same set of restriction enzymes and carrying mutated p51 domain in MCS-1. Hereafter we refer to this double mutant as RT600.

### RT139A and RT600 expression, purification, and complexes of RT/38-mer aptamer DNA

Expression and purification of RT139A containing I63C mutation was carried out as described previously (48). Briefly, the protein expression in *E. coli* BL21-CodonPlus (DE3)-RIL cells (Agilent Technologies) was induced with 1 mM IPTG and continued for 3 h at 37 °C. Cell disruption was achieved by sonication (Branson Sonifier SFX250) and the soluble fraction was subjected to immobilized metal affinity purification using a 5 ml Ni-NTA column (GE healthcare) connected to a FPLC system (GE healthcare). The 6xHis tag was cleaved from the N-terminus of p51 using a 1:10 mass ratio of HRV14 3C protease by overnight treatment on ice. The protein was further purified by anion-exchange chromatography on a Mono-Q column (GE healthcare). Expression and purification of RT600 was carried out in a similar way as with RT139A except that the 6x-His tag on p51 N-terminus was not cleaved. The Mono-Q purified RTs were stored in a buffer containing 10 mM Tris-HCl pH 8.0, 75 mM NaCl at −80 °C prior to further use. We used two 38-mer DNA-hairpin aptamers in this study, 5’-TAATTCCCCCCCTTCGGTGCTTTGCACCGAAGGGGGGG-3’ (T overhang) and 5’-TAATACCCCCCCTTCGGTGCTTTGCACCGAAGGGGGGG-3’ (A overhang), to complex with RT139 and RT600, respectively. The aptamers were purchased from Integrated DNA Technologies (Leuven, Belgium). RT/DNA-apt complexes were prepared following earlier described protocol (21) (23). Briefly, the aptamer pellet was resuspended in a buffer containing 10 mM Tris pH 8.0, 50 mM NaCl, and 1 mM EDTA, added to the protein in 1.2:1 molar ratio. The individual complexes were incubated over ice for 1 hour and purified by size-exclusion chromatography (SEC) using a Superdex 200 10/300 GL column (GE healthcare) pre-equilibrated in a buffer containing 10 mM Tris-HCl pH 8.0, 75 mM NaCl. The RT139A/DNA-apt and RT600/DNA-apt complex formation were confirmed by OD_260_:OD_280_ measurement of ∼0.95 and shift in the SEC elution profile.

### Biochemical assay

The RT inhibition assay was carried out using a Cy5-flurophore-labeled 17-mer primer (5ʹ-Cy5-CAGGAAACAGCTATGAC-3ʹ) and a template (5ʹ-GGGTGTCATAGCTGTTTCCTG-3ʹ) following a published protocol. (47) Each 20 μL enzymatic reaction mixture containing 125 nM primer-template prepared in 50 mM Tris−HCl pH 8.3, 3 mM MgCl2, 10 mM DTT, 1 μM dATP. The NNRTIs RPV, DOR and NVP were used at indicated concentrations (SI Fig. 4), and 0.006 μg of RT was added per 1 μL reaction. The primer and template were pre-annealed at 1:2 molar ratio by heating up at 95 °C and cooling down to room temperature. The reaction mixture without dATP was preincubated at 37 °C for 20 min. The reaction was initiated by adding the dATP and quenched after 1 min by adding a double volume of quenching buffer (90 % formamide, 50 mM EDTA, and 0.05 % orange G) and heating at 95 °C for 5 min. The samples were separated on a 1 mm 15 % denaturing polyacrylamide gel, and gel bands were visualized using the Typhoon FLA 9500 imaging system (GE Healthcare). The images were processed using ImageQuant TL v8.1.0.0 (GE Healthcare).

### PicoGreen Reverse transcriptase assay

The RT assay was performed with the EnzCheck Reverse Transcriptase Assay kit (Molecular Probes, Invitrogen), as described by the manufacturer. The assay is based on the dsDNA quantitation reagent PicoGreen. This reagent shows a pronounced increase in fluorescence signal upon binding to dsDNA or RNA/DNA heteroduplexes. Single-stranded nucleic acids generate only minor fluorescence signal enhancement when a sufficiently high dye: base pair ratio is applied. (49) This condition is met in the assay. Five µl of a 1 mg/ml poly(rA) template of approximately 350 bases long was annealed with 5 µl of a 50 µg/ml oligo(dT)16 primer in a molar ratio of 1:1.2 (60 min. at room temperature). The annealed template/primer was then diluted 180-fold in polymerization buffer (60 mM TrisHCl, 60 mM KCl, 8 mM MgCl_2_, 13 mM DTT, 100 µM dTTP, pH 8.1) and 17 µl of this RNA/DNA was brought into each well of a 96-well flat-black plate. To test the RT inhibition, 3 µl of compound in DMSO was added to each well before the addition of RT enzyme solution. Control wells without compound contained the same amount of DMSO. Five µl of RT enzyme solution diluted to a suitable concentration in the enzyme dilution buffer (50 mM TrisHCl, 20% glycerol, 1 mM DTT, pH 7.5) was added. The reactions were incubated at 25°C for 40 minutes and then stopped by adding 2 µl 200 mM EDTA. Heteroduplexes were then detected by addition of 173 µl 0.73 µM PicoGreen. Signals were read using an excitation wavelength of 490 nm and emission detection at 523 nm in a spectrofluorometer (Safire 2, Tecan). The results (summarized in Table 1 and detailed in SI Table. 2) are expressed as relative fluorescence i.e., the fluorescence signal of the reaction mix with compound divided by the signal of the same reaction mix without compound.

### Inhibitor complex preparation for Cryo-EM

Inhibitor stocks (NVP, RPV, and DOR) were prepared in 100% DMSO and diluted down to ∼0.5-1 mM working stocks in buffer containing 10 mM Tris-HCl pH 8.0, 75 mM NaCl immediately prior to use. Protein samples stored in 10 mM Tris-HCl pH 8.0, 75 mM NaCl at −80 °C were thawed on ice, diluted down to 0.3 mg/ml (2.3 μM) and the inhibitors were added at a 1:1.2 protein to inhibitor molar ratio. The mixtures were allowed to incubate over ice for 1 h to ascertain the formation of the respective complexes and each complex spun down in a tabletop centrifuge at 16,000 x g for 10 mins that settles any aggregates to the bottom; the sample on top was used for preparing cryo-EM grids. To find if dNTP can bind the RT/DNA/DOR complex, ∼100x molar excess dATP was added to the 1h-incubated sample of the RT/DNA/DOR complex.

### Cryo-EM grid preparation, data collection and processing

The cryo-EM grids of RT139A/DNA-apt and RT600/DNA-apt in complexes with RPV, NVP, and DOR were prepared on Quantifoil R1.2/1.3 gold mesh grids with either holey carbon or UltrAuFoil films (Quantifoil, Germany). The holey carbon film grids were pre-cleaned in chloroform for 2-3 h at room temperature and allowed to dry overnight prior to use while the ones with UltrAuFoil film were used without any cleaning. The grids were glow-discharged for 45 sec at 25 mA with the chamber pressure set at 0.3 mBar (PELCO easiGlow; Ted Pella). The grids were mounted in the sample chamber of a Leica EM GP set at 8 ^0^C and 95% relative humidity. Optimized grids were obtained by applying 3 μl of the sample at 0.3 mg/ml, incubating for 30 sec, back-blotting for 14 sec using Whatman Grade 1 filter paper, and plunge-freezing in liquid ethane at −172 °C. The prepared grids were then clipped and mounted on a 200 kV Glacios TEM (ThermoFisher Scientific) equipped with autoloader and Falcon 3 direct electron detector as installed in our laboratory. Cryo-EM data sets were collected from the frozen hydrated samples in counting mode on the Glacios TEM using EPU software version 2.9.0 (ThermoFisher Scientific). The movies were collected at a nominal magnification of 150,000x yielding a pixel size of 0.97 Å. Each movie was collected with 40 frames, where each frame received ∼1 e/Å^2^, for a total dose of 40 e/Å^2^ and subsequently written as a gain-corrected MRC file. Data collection statistics are listed in Supplementary Table S1.

### Cryo-EM data processing

Individual movie frames were motion-corrected and aligned using MotionCor2 (50) as implemented in the Relion-3.1 package (51) and the contrast transfer function (CTF) parameters were estimated by CTFFIND-4 (52). The particles were automatically picked using the reference-free Laplacian-of-Gaussian routine in Relion-3.1. The picked particles were initially classified as 3D classes which proved to be more efficient than initial 2D classification. The 3-D classification generated a distinct single class of particles with meaningful map connectivity and highest estimated resolution compared to other classes. The particles in the good 3D class were re-extracted and subjected to 2D and 3D classifications. The final set of particles for each RT/DNA/NNRTI complex was used to calculate gold-standard auto-refined maps, which were further improved by B-polishing and CTF-refinement. No additional stable classes representing states such as RT/DNA or RT/NNRTI were observed. All data processing was carried out using Relion-3.1.

### Model building

RT/DNA-apt crystal structure (PDB Id. 5HP1) was used as the starting model for fitting the atomic structure to the density map of RT/DNA-apt/NVP complex. The RT/DNA-apt/NVP structure was subsequently used as the template for building other structures reported in this study. Manual model fitting to the density map was carried out in COOT (53) followed by real-space model refinement using Phenix 1.19 (54). The inhibitor model and restrain files were obtained from PDB. All structure figures were prepared with PyMOL (https://pymol.org/2/) and Chimera (55).

## Data Availability

The density maps and atomic coordinates for the structures reported in this paper have been deposited in the Electron Microscopy Databank (EMDB) and Protein Data Bank (PDB), respectively. The accession codes are: EMD-14457 and 7Z24 for wild-type RT/DNA/NVP; EMD-14458 and 7Z29 for E138K/M184I mutant RT/DNA/NVP; EMD-14462 and 7Z2D for wild-type RT/DNA/RPV; EMD-14463 and 7Z2E for E138K/M184I mutant RT/DNA/RPV; EMD-14465 and 7Z2G for wild-type RT/DNA/DOR; EMD-14466 and 7Z2H for E138K/M184I mutant RT/DNA/DOR.

## Acknowledgments

This paper is dedicated to the memory of our dear colleague Sergio E. Martinez, who passed away recently. Over the years, Sergio contributed to studies of macromolecule structures and structure-based drug design including the research published in this manuscript. The study was supported by Rega Virology and Chemotherapy internal grants to K.D., and E.A. acknowledges National Institutes of Health (NIH) R01 AI027690 for support.

