## Supplementary material for "Cryo-EM structures of Doravirine and Rilpivirine with HIV-1 Reverse Transcriptase/DNA Aptamer – Nonnucleoside Inhibitor Resistance by E138K and M184I Mutations": SI Table 1, SI Table 2, SI Fig. 1, SI Fig. 2, SI Fig. 3, SI Fig. 4, SI Fig. 5

**SI Table 1: Single particle cryo-EM data and structure analysis statistics.**

| | Wt RT/ DNA/**NVP** | wt RT/ DNA/**RPV** | wt RT/ DNA/**DOR** | M184I + E138K RT/ DNA/**NVP** | M184I + E138K RT/ DNA/**RPV** | M184I + E138K RT/ DNA/**DOR** |
|---|---|---|---|---|---|---|
| PDB Id/EMBD Id | 7Z24/EMD -14457 | 7Z2D/EMD -14462 | 7Z2G/EMD-14465 | 7Z29/EMD -14458 | 7Z2E/EMD -14463 | 7Z2H/EMD-14466 |
| | **Data collection** | | | | | |
| Grid type; Number of grids | Quantifoil Au R1.2/1.3; 1 | Quantifoil Au R1.2/1.3; 1 | Quantifoil Ultra Au R1.2/1.3; 1 | Quantifoil Au R1.2/1.3; 1 | Quantifoil Au R1.2/1.3; 1 | Quantifoil Ultra Au R1.2/1.3; 1 |
| Microscope; Voltage (kV); Detector | Glacios; 200kV; Falcon 3 | Glacios; 200kV; Falcon 3 | Glacios; 200kV; Falcon 3 | Glacios; 200kV; Falcon 3 | Glacios; 200kV; Falcon 3 | Glacios; 200kV; Falcon 3 |
| Magnification; Pixel size (Å) | 150,000x; 0.97 | 150,000x; 0.97 | 150,000x; 0.97 | 150,000x; 0.97 | 150,000x; 0.97 | 150,000x; 0.97 |
| Dose ($e^-/Å^2$/frame); Number of frames per movie | 1.0; 40 | 1.0; 40 | 1.0; 40 | 1.0; 40 | 1.0; 40 | 1.0; 40 |
| Total dose ($e/Å^2$); | 40 | 40 | 40 | 40 | 40 | 40 |
| Total exposure time (sec) | 55 | 55 | 55 | 55 | 55 | 55 |
| Number of micrographs recorded | 1,938 | 1,647 | 1,352 | 1,981 | 1,850 | 2,104 |
| Defocus range (Å) | 8,000 – 20,000 | 8,000 – 20,000 | 8,000 – 20,000 | 8,000 – 20,000 | 8,000 – 20,000 | 8,000 – 20,000 |
| | **Data processing** | | | | | |
| Number of micrographs used | 1,910 | 1,451 | 1,299 | 1,354 | 1,769 | 1,067 |
| Number of particles picked | 1,792,933 | 997,218 | 1,235,740 | 1,323,229 | 1,702,421 | 970,781 |
| Particles used in final map | 298,193 | 200,331 | 178,579 | 304,552 | 259,849 | 148,135 |
| Map resolution (FSC 0.143; Å) | 3.32 | 3.38 | 3.65 | 3.38 | 3.45 | 3.58 |
| Map sharpening B factor ($Å^2$) | -150.4 | -158.3 | -194.3 | -163.4 | -166.4 | -140.1 |
| | **Model fitting** | | | | | |
| Total number of atoms | 8,303 | 8,591 | 8,406 | 8,307 | 8,479 | 8,414 |
| Expt. map/model correlation | 0.70 | 0.69 | 0.76 | 0.71 | 0.68 | 0.61 |
| Map/ligand correlation | 0.82 | 0.80 | 0.76 | 0.83 | 0.77 | 0.67 |
| Average B factor ($Å^2$) - Protein | 28.05 | 29.85 | 31.70 | 30.05 | 24.00 | 45.98 |
| - DNA | 86.46 | 64.22 | 56.87 | 74.55 | 51.76 | 73.98 |

| | | | | | | |
|---|---|---|---|---|---|---|
| - Ligand | 25.79 | 26.52 | 13.81 | 25.81 | 8.03 | 26.39 |
| Clash score | 7.61 | 7.33 | 7.71 | 7.43 | 8.08 | 8.84 |
| Ramachandran favored/outlier | 97.1/0.0 | 96.1/0.0 | 96.1/0.0 | 96.8/0.1 | 96.7/0.1 | 95.5/0.1 |

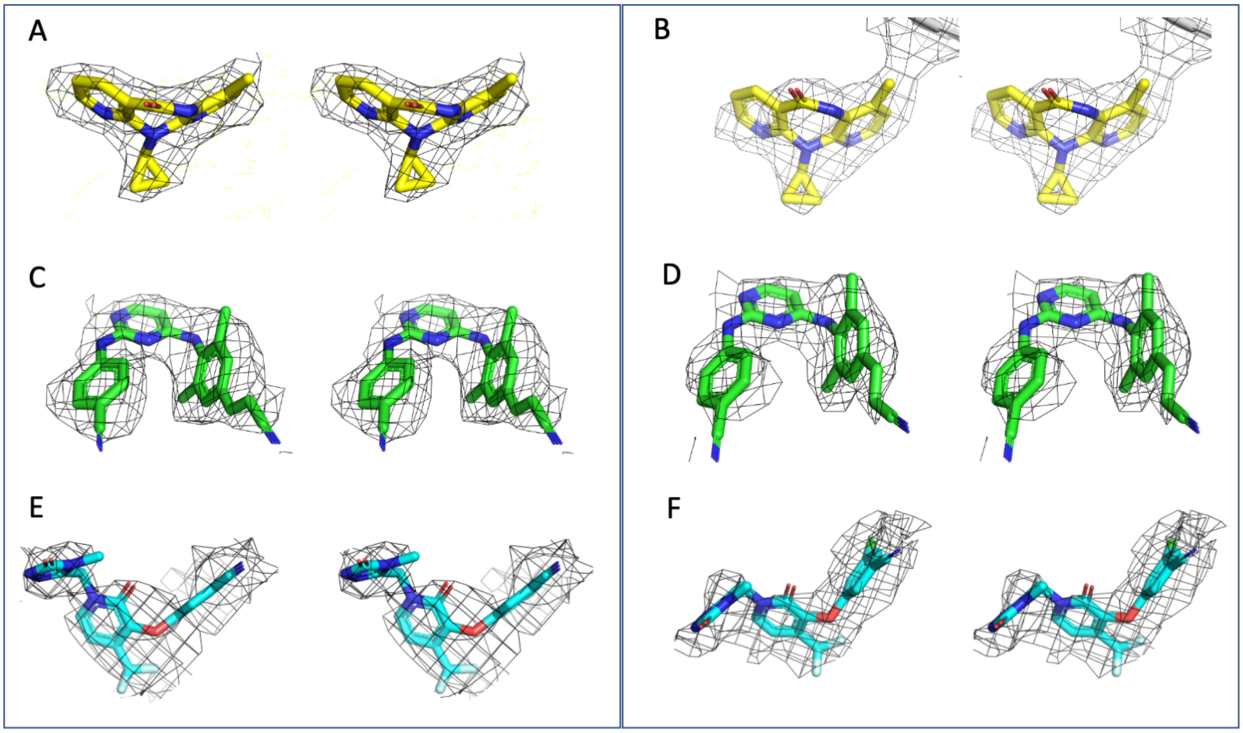

**SI Figure 1. Wall-eyed stereo views of fitting of NNRTIs to single-particle cryo-EM density maps.** **A.** The density for NVP in the wild-type RT/DNA/NVP ternary complex displayed at contour level 4.0σ. **B.** The density for NVP in the E138K/M184I mutant RT/DNA/NVP ternary complex displayed at contour level 3.5σ. **C.** The density for RPV in the wild-type RT/DNA/RPV ternary complex displayed at contour level 3.2σ. **D.** The density for RPV in the E138K/M184I mutant RT/DNA/RPV ternary complex displayed at contour level 3.0σ. **E.** The density for DOR in the wild-type RT/DNA/DOR ternary complex displayed at contour level 2.2σ. **F.** The density for DOR in the E138K/M184I mutant RT/DNA/DOR ternary complex displayed at contour level 2.2σ.

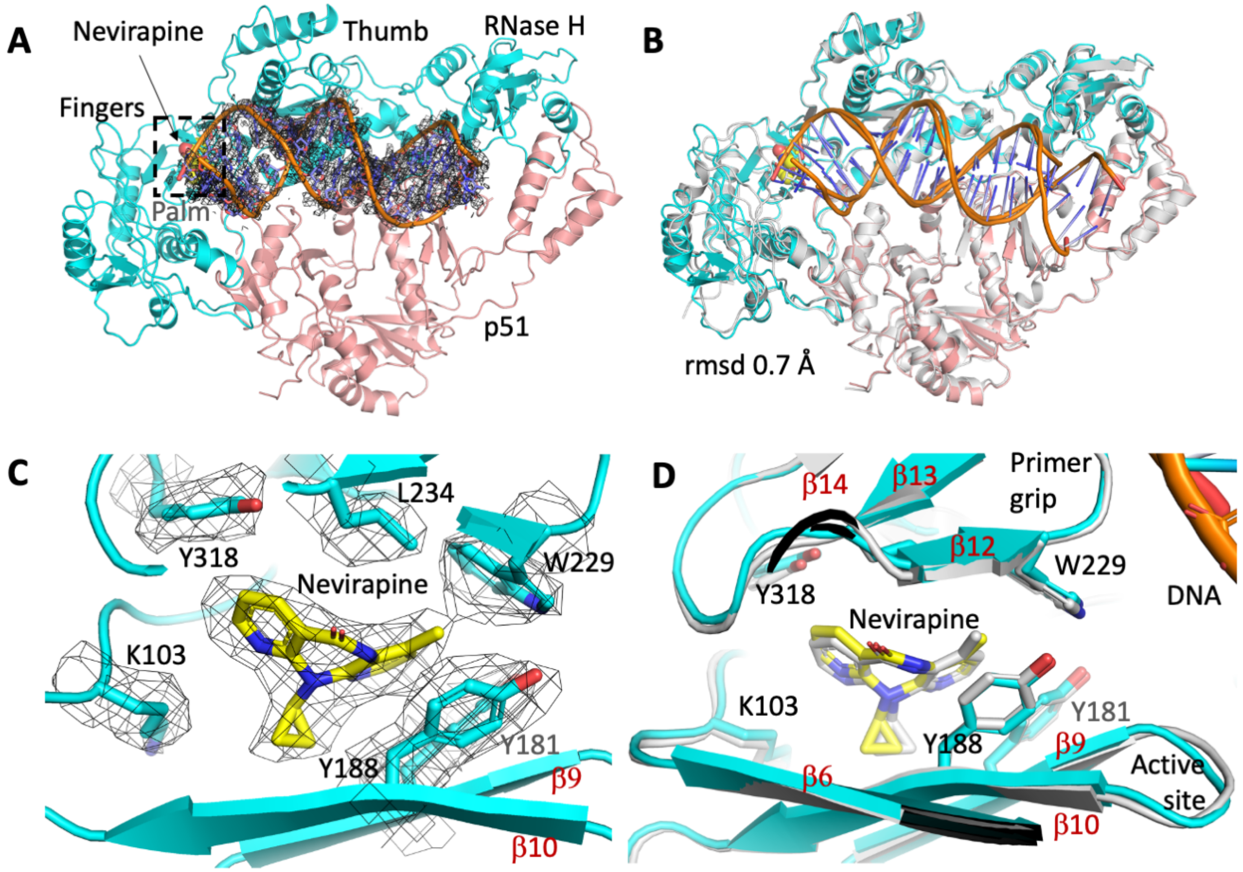

**SI Figure 2. Cryo-EM structure of wild-type HIV-1 RT/DNA-apt/NVP and its comparison with the previously reported crystal structure of RT/DNA/NVP complex** (PDB Id. 3V81). **A.** Cryo-EM structure of wild-type HIV-1 RT/DNA-apt/NVP complex. The p66 and p51 subunits are colored cyan and salmon, respectively; the DNA aptamer is modeled into the density map. **B**. Superposition of the crystal structure (gray) on the cryo-EM structure. **C.** A zoomed view of the RT/DNA/NVP structure at the NNIBP region; the cryo-EM density for NVP and surrounding residues are countered at 4σ. D. The zoomed view showing the NNIBP region of the superimposed structures in panel B.

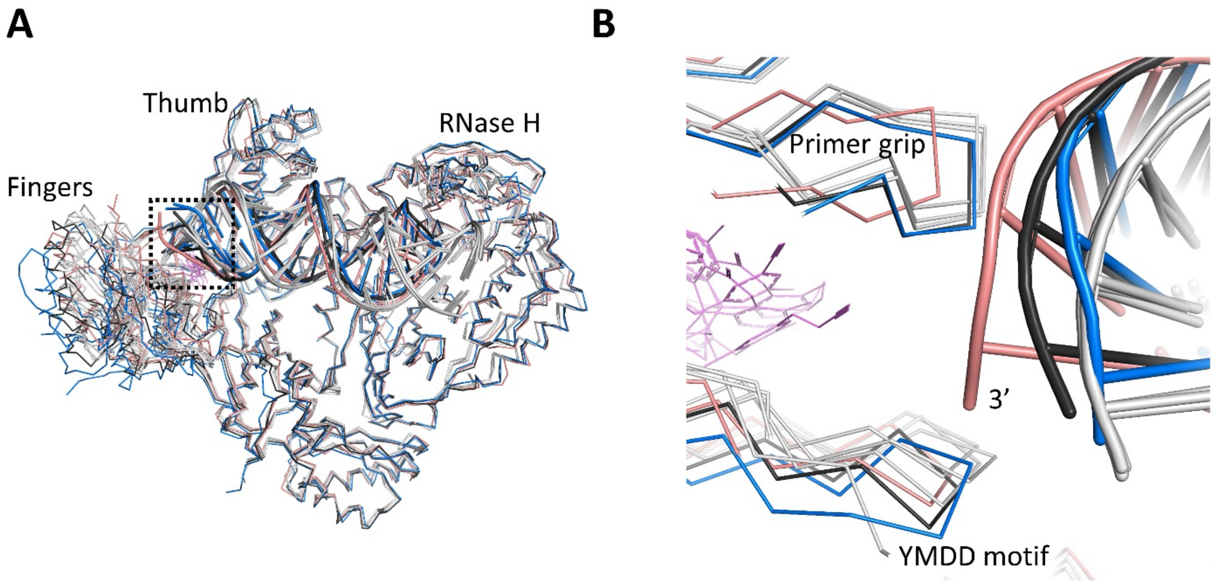

**SI Figure 3. Overlay of RT/dsDNA/NNRTI and RT/dsRNA/NNRTI structures. A.** C$\alpha$ superposition of wild-type RT/DNA/DOR (salmon), RT/DNA/RPV (blue), RT/DNA/NVP (black), RT/dsRNA (PDB Id. 7KJV), RT/dsRNA/NVP (PDB Id. 7KJX), and RT/dsRNA/EFV (PDB Id. 7KJW); all RT/dsRNA structures are in gray. B. A zoomed view of at the active site of the superimposed structures. The primer grip and the nucleic acid track of DOR complex and the YMDD motif of RPV complex deviate the most.

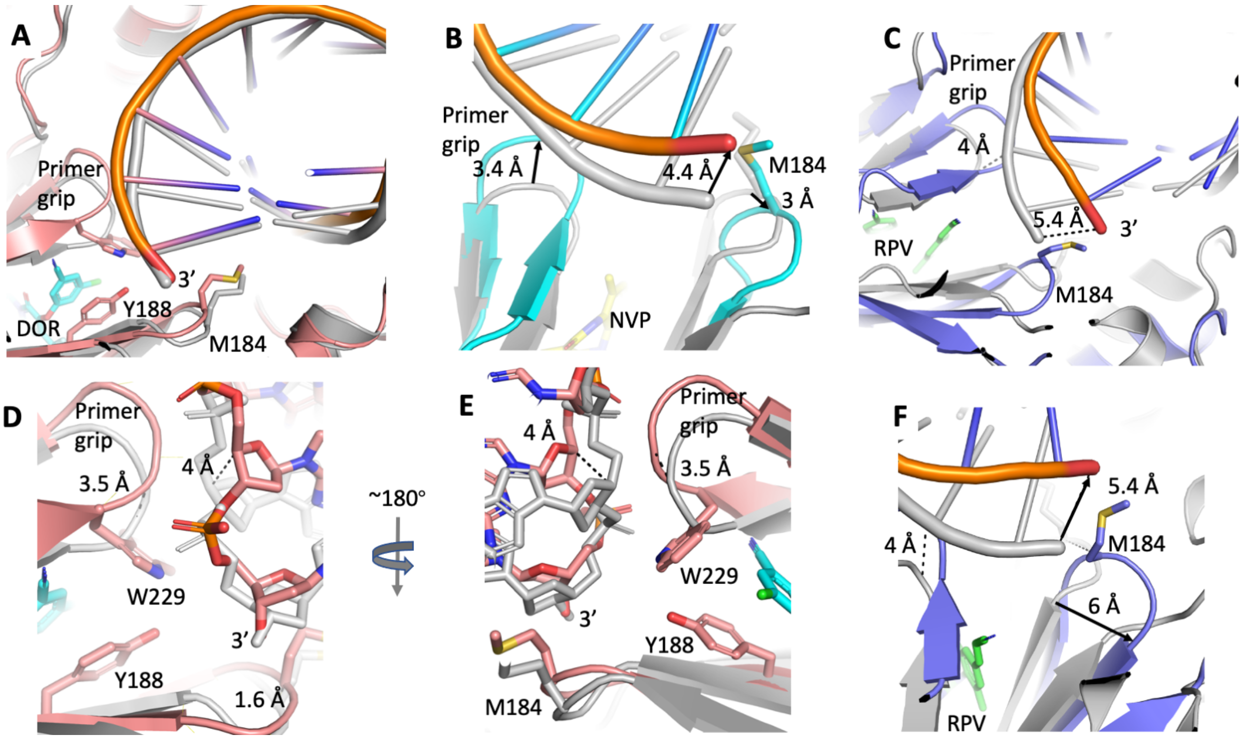

**SI Figure 4. Impacts of different NNRTIs on catalytically active RT/DNA conformation.** C$\alpha$ superpositions of DOR (**A**), NVP (**B**), and RPV (**C**) on RT/DNA structure (PDB Id. 5D3G); the NNRTI ternary complexes are colored and the RT/DNA binary complex in all panels is in gray. For DOR complex 840 Ca, for NVP complex 770 Ca, and 786 Ca for RPV complex are aligned with RT/DNA complex with RMSD of 0.92, 1.5, and 1.7 Å, respectively. **D** and **E** show two views of the active site region of the aligned RT/DNA/DOR and RT/DNA structures. The primer 3' end in DOR complex is positioned near the polymerase active site as in RT/DNA complex, however, the primer P-1 nucleotide has moved by ~4 Å along with the primer in the DOR complex when compared with the RT/DNA structure. **F.** Active-site region in the RPV complex shows that the YMDD loop has shifted to right by ~6 Å compared to that in RT/DNA complex.

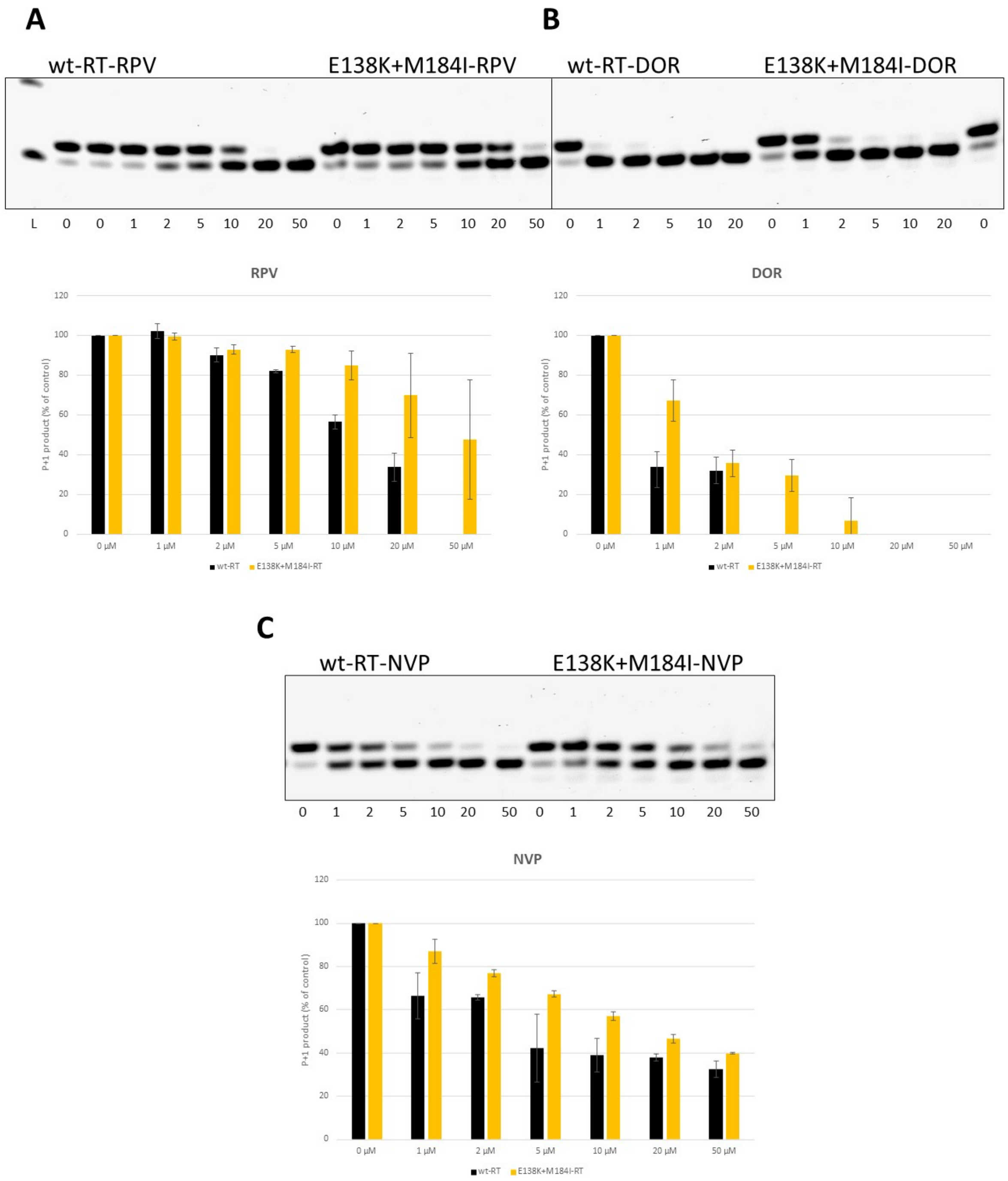

**SI Figure 5. Single-nucleotide incorporation HIV-1 RT polymerization assay**. A set of dose dependent HIV-1 RT inhibition assay was carried out using a Cy5-flurophore-labeled 17-mer primer and a 21-mer template for (**A**) RPV, (**B**) DOR and (**C**) NVP. Each 20 µL reaction mixture contained 0.006 µg of either wild-type or E138K/M184I mutant RT per 1 µL reaction, 125 nM primer-template complex prepared in 50 mM Tris−HCl pH 8.3, 3 mM MgCl2, 10 mM DTT, 1 µM dATP, and the NNRTIs at indicated concentrations. The bar charts represent percentage of product formed (Y-axis) at different inhibitor concentration (X-axis) by wt (black) and E138K/M184I (yellow) RT.

**SI Table 2: Activity of NVP, RPV and DOR against wt- and E138K/M184I-RT and $IC_{50}$ determination.** Activity assay was performed in duplicates using EnzChek™ Reverse Transcriptase Assay Kit (ThermoFisher Scientific).

| **wt-RT (µg/ml)** | 100 | 20 | 4 | 0.8 | 0.16 | 0.032 | 0.0064 | 0.00128 | **$IC_{50}$** | **av. $IC_{50}$** | SD |
|---|---|---|---|---|---|---|---|---|---|---|---|
| Nevirapine (1) | 0.06 | 0.12 | 0.17 | 0.25 | 0.46 | 0.70 | 0.90 | 0.98 | 0.12 | | |
| Nevirapine (2) | 0.07 | 0.13 | 0.21 | 0.28 | 0.47 | 0.72 | 0.88 | 0.98 | 0.13 | 0.13 | 0.01 |
| **wt-RT (µg/ml)** | 10 | 2 | 0.4 | 0.08 | 0.016 | 0.0032 | 0.00064 | 0.000128 | **$IC_{50}$** | **av. $IC_{50}$** | SD |
| Rilpivirine (1) | -0.01 | 0.01 | 0.05 | 0.08 | 0.21 | 0.52 | 0.84 | 0.99 | 0.0036 | | |
| Rilpivirine (2) | -0.01 | 0.01 | 0.04 | 0.08 | 0.21 | 0.56 | 0.89 | 0.98 | 0.0042 | 0.0039 | 0.0004 |
| **wt-RT (µg/ml)** | 2 | 0.4 | 0.08 | 0.016 | 0.0032 | 0.00064 | 0.000128 | 0.000026 | **$IC_{50}$** | **av. $IC_{50}$** | SD |
| Doravirine (1) | 0.04 | 0.09 | 0.18 | 0.40 | 0.75 | 0.90 | 0.91 | 1.02 | 0.010 | | |
| Doravirine (2) | 0.05 | 0.09 | 0.18 | 0.41 | 0.84 | 0.96 | 1.02 | 1.01 | 0.011 | 0.011 | 0.001 |
| **E138K+M184I-RT (µg/ml)** | 100 | 20 | 4 | 0.8 | 0.16 | 0.032 | 0.0064 | 0.00128 | **$IC_{50}$** | **av. $IC_{50}$** | SD |
| Nevirapine (1) | 0.04 | 0.10 | 0.14 | 0.23 | 0.38 | 0.64 | 0.83 | 0.85 | 0.076 | | |
| Nevirapine (2) | 0.04 | 0.10 | 0.14 | 0.24 | 0.43 | 0.67 | 0.88 | 0.97 | 0.10 | 0.088 | 0.017 |
| **E138K+M184I-RT (µg/ml)** | 10 | 2 | 0.4 | 0.08 | 0.016 | 0.0032 | 0.00064 | 0.000128 | **$IC_{50}$** | **av. $IC_{50}$** | SD |
| Rilpivirine (1) | -0.01 | 0.03 | 0.08 | 0.16 | 0.37 | 0.73 | 0.91 | 0.98 | 0.0089 | | |
| Rilpivirine (2) | -0.01 | 0.03 | 0.08 | 0.16 | 0.39 | 0.69 | 0.95 | 0.99 | 0.0089 | 0.0089 | 0.0000 |
| **E138K+M184I-RT (µg/ml)** | 2 | 0.4 | 0.08 | 0.016 | 0.0032 | 0.00064 | 0.000128 | 0.000026 | **$IC_{50}$** | **av. $IC_{50}$** | SD |
| Doravirine (1) | 0.06 | 0.17 | 0.34 | 0.56 | 0.86 | 0.97 | 0.95 | 0.97 | 0.025 | | |
| Doravirine (2) | 0.06 | 0.14 | 0.34 | 0.57 | 0.92 | 0.97 | 1.04 | 1.01 | 0.026 | 0.026 | 0.001 |
